# A simple and highly efficient method for multi-allelic CRISPR-Cas9 editing in primary cell cultures

**DOI:** 10.1101/2020.07.01.183145

**Authors:** Pia Hoellerbauer, Megan Kufeld, Sonali Arora, Hua-Jun Wu, Heather M. Feldman, Patrick J. Paddison

## Abstract

**Background:** CRISPR-Cas9-based technologies have revolutionized experimental manipulation of mammalian genomes. None-the-less, limitations of the delivery and efficacy of these technologies restrict their application in primary cells.

**Aims:** To create an optimized protocol for penetrant, reproducible, and fast targeted insertion-deletion mutation (indel) formation in cell cultures derived from primary cells, using patient-derived glioblastoma (GBM) stem-like cells (GSCs) and human neural stem/progenitor cells (NSCs) for proof-of-concept experiments.

**Methods:** We employed transient nucleofection of Cas9:sgRNA ribonucleoprotein complexes using chemically synthesized 2’-O-methyl 3’phosphorothioate-modified sgRNAs and purified Cas9 protein. Indel frequency and size distribution were measured via computational deconvolution of Sanger sequencing trace data. Western blotting was used to evaluate protein loss. RNA-seq in edited NSCs was used to assess gene expression changes resulting from knockout of tumor suppressors commonly altered in GBM.

**Results:** We found that with this optimized technique, we can routinely achieve >90% indel formation in only 3 days, without the need to create clonal lines for simple loss-of-function experiments. We observed near-total protein loss of target genes in cell pools. Additionally, we found that this approach allows for the creation of targeted genomic deletions. We also demonstrated the utility of this method for quickly creating a series of gene knockouts that allow for the study of oncogenic activities.

**Conclusion:** Our data suggest that this relatively simple method can be used for highly efficient and fast gene knockout, as well as for targeted genomic deletions, even in hyperdiploid cells (such as GSCs). This represents an extremely useful tool for the cancer research community when wishing to inactivate not only coding genes, but also non-coding RNAs, UTRs, enhancers, and promoters. This method can be readily applied to diverse cell types by varying the nucleofection conditions.

## BACKGROUND

In bacteria and archaea, the CRISPR-Cas (Clustered, Regularly Interspaced, Short Palindromic Repeats (CRISPR)–CRISPR-associated (Cas)) pathway acts as an adaptive immune system, conferring resistance to genetic parasites and bacteriophage (1, 2). CRISPR-Cas systems are able to target and degrade DNA (1, 2), and this property has been harnessed for directed genome editing in prokaryotes and (more recently) eukaryotes, including human cells (3-6), using the type II CRISPR-Cas system from *Streptococcus pyogenes*. In its simplest form, this system consists of a complex of two components, the Cas9 protein and an sgRNA. Cas9 is an RNA–guided DNA endonuclease. The sgRNA is a chimeric guide RNA composed of a ~20nt ‘protospacer’ sequence, which is used for target recognition, and a structural RNA required for Cas9:sgRNA complex formation (i.e. tracrRNA). In addition, DNA cleavage by Cas9 occurs only in the presence of an appropriate protospacer adjacent motif (PAM) at the 3’ end of the protospacer sequence in the target genomic locus (for Cas9 this is “NGG”, where N is any nucleotide (2)).

When Cas9 and an sgRNA are expressed together, a double-strand DNA (dsDNA) break is created about 3 bp upstream of the PAM site (7, 8). This break is then repaired by the cell either via the high-fidelity homology-directed repair (HDR) pathway, or much more commonly, via the error-prone non-homologous end joining (NHEJ) pathway, which leaves behind repair scars in the form of small insertion-deletion (indel) mutations (9, 10). When these indels occur in an exon, they can cause frameshifts and premature stop codons in the target gene, effectively ablating protein function (4, 6, 11).

Cas9:sgRNA targeting efficiency in human cells varies considerably depending on the methods, reagents, and cell types used. In general, successful generation of indels using transient DNA transfection occurs in a range of ~1-30% (8). However, it was shown that lentiviral-based stable expression of Cas9:sgRNA greatly improves targeting efficiency to >90% (2, 12, 13). As a direct result, we and others have successfully performed pooled lentiviral-based sgRNA screens in various human cell types (12-14). However, retesting single sgRNAs from these screens, especially those targeting essential genes, can prove challenging. For example, in human neural stem/progenitor cells (NSCs) and patient-derived glioblastoma stemlike cells (GSCs), we have observed that “all-in-one” lentiviral-based CRISPR systems can result in protracted windows of indel formation and phenotypically mixed populations, requiring incubation of up to 12 days to achieve complete indel formation (14). As a result, it can be difficult to set up rigorous experiments analyzing a particular gene knockout (KO) when cell populations contain variable mixtures of wild-type (wt) and indel-containing alleles and, if the target gene is essential, thereby contain mixtures of alive and dead cells. This represents a critical experimental limitation of the use of CRISPR-Cas9 platforms in primary cells. As a result, we wished to create an optimized protocol that would allow maximal targeted indel formation over the shortest possible experimental window.

We found a robust method that utilizes transient nucleofection of *in vitro-formed* Cas9:sgRNA ribonucleoprotein (RNP) complexes using chemically synthesized 2’-O-methyl 3’phosphorothioate-modified sgRNAs and purified Cas9 protein (Fig. 1A). With this optimized approach, we are able to achieve >90% indel formation in multiple human GSCs and NSCs in only 3 days, without the need to create clonal lines for simple loss-of-function experiments. In addition, we find that this approach allows for the creation of targeted deletions in cell pools or cell clones, depending on the size of the desired deletion. Here we present these results illustrating the utility of this method and a detailed step-by-step protocol (Supplementary Materials).

**Figure 1.**
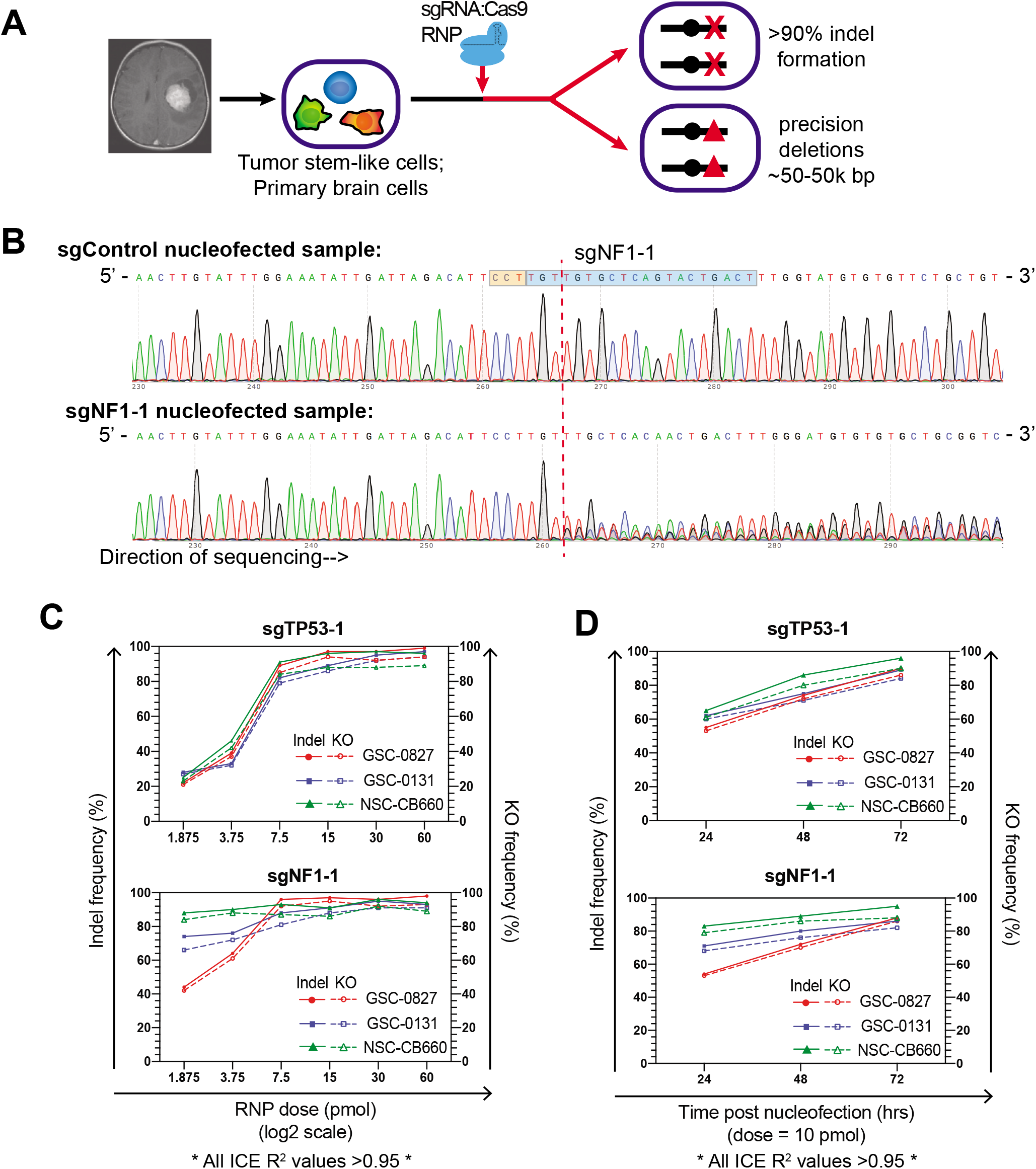
Highly efficient and fast indel formation using RNPs composed of purified Cas9 and chemically synthesized 2’-O-methyl 3’phosphorothioate-modified sgRNA. (A) Overview of CRISPR RNP targeting strategy. (B) Sanger trace example for a pool of NSC-CB660 nucleofected with 15 pmol RNPs using either a non-targeting control sgRNA or sgNF1-1 (72 hours post-nucleofection). Sequence was created using a reverse sequencing primer. Blue shaded box denotes sgRNA, orange shaded box denotes PAM sequence, and dotted red line represents sgRNA cut site. The sgNF1-1 treated cells produce a clean sequence trace that mirrors the control up until the sgRNA cut site, at which point the trace begins to represent the compounding effect of multiple overlapping traces due to various indel mutations. Sanger trace data like this was used in conjunction with a freely-available bioinformatics tool (ICE) in order to predict CRISPR editing sequence distribution in cell pools. (C) CRISPR editing efficiency as a function of RNP dose for 2 different sgRNAs. Solid data lines denote indel frequency while dotted lines denote predicted KO frequency (% of predicted sequences that result in a frameshift or an indel ≥21 bp in length). All samples were harvested 72 hours post-nucleofection. (D) CRISPR indel frequency and KO frequency as a function of time post-nucleofection for 2 different sgRNAs. A dose of 10 pmol was used.

## RESULTS

### Nucleofection of Cas9:sgRNA (chemically synthesized, modified) RNPs results in highly penetrant generation of small indels in human GSCs and NSCs

Due to the previously-discussed challenges with indel formation using lentiviral-based Cas9:sgRNA delivery, we explored alternative approaches, including the use of Cas9:sgRNA ribonucleoprotein (RNP) complexes composed of purified Cas9 protein and purified gRNA. Such RNPs have recently been used effectively for several applications, including gene loss-of-function in human cell lines (15-17) and ES cells (15), editing of CXCR4 in human T cells (18), HDR tests in human cells via insertion of restriction sites (15, 19), epitope tagging in mouse NSCs (20), and studying effects of sgRNA sequence on editing efficiency (21). The efficiencies reported for *in vitro* editing in these contexts are most often in the range of ~15-60%, with most studies reporting *maximum* (not routine) efficiencies ≤80%. We wanted to further improve upon these RNP methods so we could routinely achieve high efficiency, multi-allelic editing in GSCs and NSCs (Fig. 1A).

Since most studies using RNPs utilize *in vitro*-transcribed gRNA, we instead tested chemically synthesized sgRNAs with 2’-O-methyl 3’ phosphorothioate modifications in the first and last 3 nucleotides, which are more nuclease resistant than unmodified sgRNAs and therefore likely increase RNP half-life (22, 23). We formed RNPs by combining these sgRNAs with purified sNLS-SpCas9-sNLS nuclease and then delivered them via nucleofection, a modified electroporation technique developed by Amaxa (now Lonza) that allows direct transfer of nucleic acids into the nucleus of mammalian cells in culture.

To follow indel formation in cell pools, we employed a method that uses Sanger sequencing of sgRNA target site-spanning PCR amplicons followed by computational trace decomposition of the control and experimental traces to predict indel frequency and KO frequency (24, 25), where KO frequency is the percent of predicted sequences that result in a frameshift or an indel ≥21 bp in length (25) (Fig. 1B). In order to determine the efficiency and timing of indel formation, we measured indel and KO frequency for RNP doses ranging from ~2-60 pmol (Fig. 1C) and at 24, 48, and 72 hours post-nucleofection (Fig. 1D) in diploid NSC-CB660 cells, near-diploid GSC-0131 cells, and hypertriploid GSC-0827 cells, for single sgRNAs targeting *TP53* exon 7 (sgTP53-1) and *NF1* exon 2 (sgNF1-1). *TP53* and *NF1* are both located on chromosome 17, of which GSC-0131 have 2 copies and GSC-0827 have 3 copies. These experiments revealed that high (>90%) multi-allelic indel efficiencies can be achieved starting at RNP doses of 7.5-15 pmol in GSCs and NSCs (Fig. 1C). Furthermore, we observed that 50-70% of editing has occurred by 24 hours post-nucleofection, reaching its maximum by 72 hours (Fig. 1D). Similar penetrance of sgRNAs targeting other genes was also observed in multiple GSC isolates and NSCs which had been immortalized and oncogenically transformed (Fig. S1). Because many of our sgRNA sequences had been prevalidated through our lentiviral screens (14) (e.g. sgTP53-1; sgNF1-1; 13 of 23 sgRNAs in Fig. S1), these experiments illustrate representative results for active RNPs. Of note, several of the genes targeted are present at hyperdiploid levels in GSCs; for instance, GSC-G166 contain 3 copies of *SCAP, FBXO42*, and *GMPPB*. Furthermore, targets included top scoring essential genes for both GSCs and NSCs, which cause profound viability loss (14), indicating that the high efficiencies we observe are *not* simply due to outgrowth of edited cells.

### RNP nucleofection allows for targeted deletion of several hundred bp genomic regions

In our efforts to assess RNP efficiency and dosing, we also observed that for sgTP53-1 and sgNF1-1, the KO efficiency closely mirrored the indel efficiency (Fig. 1C-D). In these cases, this is due to repair bias at the target sites, where each sgRNA produced a high percentage of 1bp insertions in GSCs and a mixture of frameshifting 1bp insertions/deletions and 2 bp deletions in NSCs (Fig. 2A-B, top panels). This is consistent with a recent analysis of sgRNA targeting repair events in human cells, which found that frameshift frequencies are higher than expected (81% versus the expected 67% for random NHEJ-mediated repair) due to unexpectedly high 1 bp insertion/deletion events (11).

**Figure 2.**
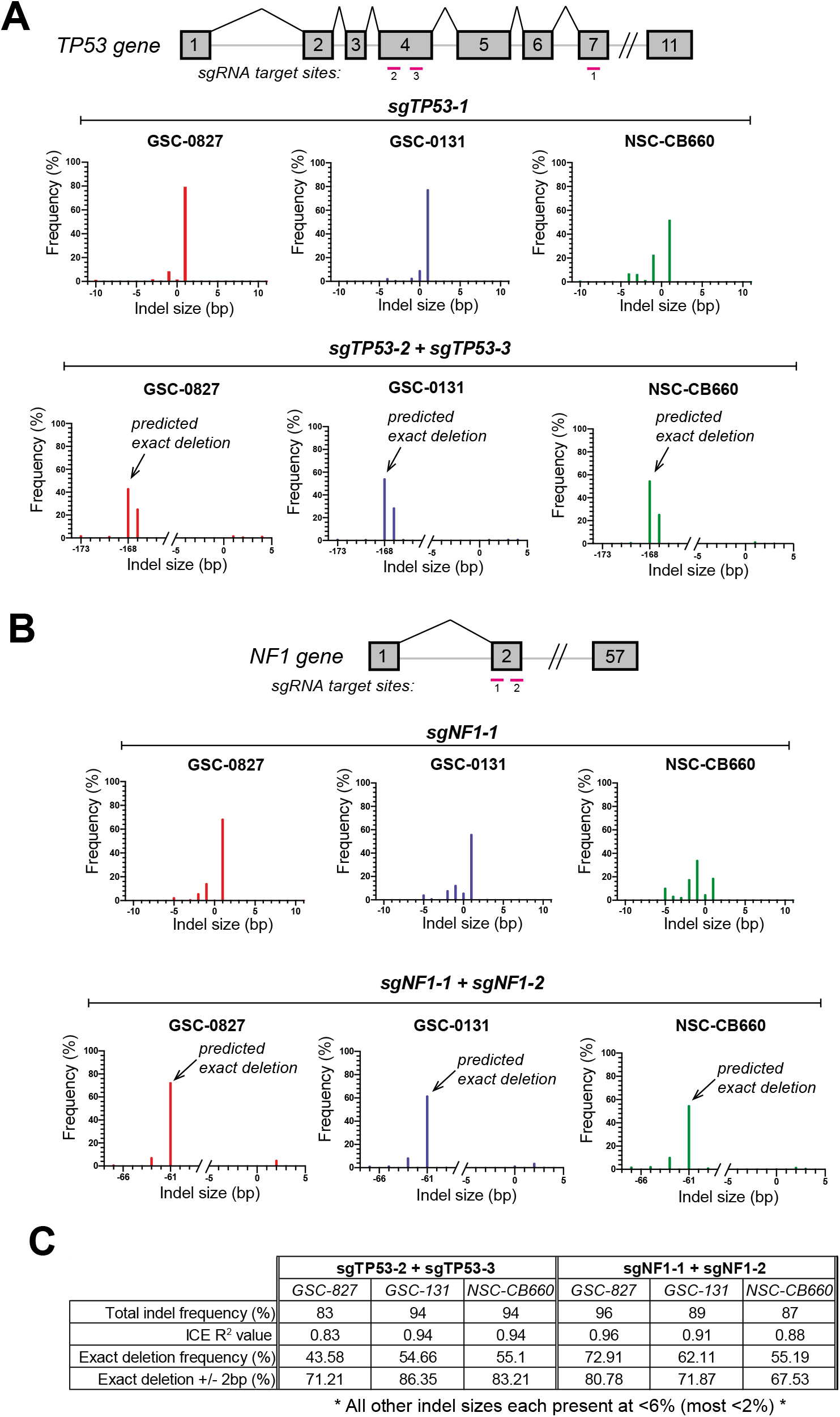
Cas9:sgRNA RNPs can be used to create small targeted deletions. (A) Indel size distribution of predicted indel sequences for cell isolates nucleofected with individual sgTP53s (top panel) or two simultaneous sgTP53s (bottom panel) (168 bp apart). (B) Indel size distribution of predicted indel sequences for cell isolates nucleofected with individual sgNF1s (top panel) or two simultaneous sgNF1s (bottom panel) (61 bp apart). (C) Summary of dual sgRNA indel profiles shown in (A) and (B).

Given the precise and reproducible nature of the indels created for sgNF1-1 and sgTP53-1, we wondered if using two sgRNAs in close proximity (e.g. 50-1000 bp) would favor the production of precise deletions using our protocol. This was indeed the case when we simultaneously nucleofected with two sgRNAs targeting *TP53* or *NF1* (Fig. 2A-B, bottom panels). We observed remarkably high predicted exact deletion frequencies between 44-73% for GSCs and NSCs (Fig. 2C). Allowing +/-2bp for the deletion size window further increased the predicted “near-precise” deletion efficiencies to 68-86% (Fig. 2C). It is also important to note that in this multiple sgRNAs scenario, the bioinformatic predictions for total indel frequencies were somewhat reduced due to adjustment for slightly lower regression fit R^2^ values. However, essentially 0% wt sequences were predicted in the trace data (see indel size of “0” in bottom panels of Fig. 2A and 2B), suggesting that the total indel frequencies – and thereby also the deletion frequencies – may actually have been even higher. We observed similar near-precise deletion results when nucleofecting with two other sets of double sgRNAs (Fig. S2).

To further investigate these results, we designed sets of 3 sgRNAs to target 13 different non-coding genomic loci on chromosomes X, 2, 5, 13, 15, and 21 in GSC-0827 cells, which contain 4, 3, 3, 2, 3, and 3 copies of these chromosomes, respectively. These sgRNA sequences were designed using the Broad GPP Web Portal (26) and were used without prevalidation. Each target locus was defined by two outer/flanking sgRNA cut sites (176-981 bp apart) and a third sgRNA targeting roughly the midpoint (Fig. S3A). Five days after nucleofection using these sgRNA pools (compared to a non-targeting control sgRNA), deletion production was visualized via PCR using primers flanking the outermost cut sites (Fig. S3A). The results showed that deletions, spanning either the flanking cut sites or a flank-to-mid cut, were dramatically favored over simple small indels (which are contained in the ~wt-sized band due to lack of separation for size differences of only a few bp) (Fig. S3B). These results suggest that deletions that are near-precise lengths of <1000bp can be readily generated using this protocol.

### RNP nucleofection generates dramatic protein loss in cell pools

Given the potential of our approach to create highly penetrant multi-allelic KOs in pools, we wanted to further demonstrate its utility by creating a series of KO mutants in human NSCs. Previously, we and others have used ectopic expression of human oncogenes (e.g. *EGFRvIII, RasV12, myr-Akt1, CCND1, CDK4^R24C^*, dominant-negative *TP53*) to partially or completely transform NSCs (14). Our current method afforded us the opportunity to affect the same pathways by instead creating loss-of-function mutations in tumor suppressors. We chose to successively target four genes commonly mutated or deleted in adult GBM tumors: *TP53, CDKN2A, PTEN*, and *NF1*, which affect distinct pathways required for glioma progression, including the p53-pathway, the Rb-axis, the PI3-k/Akt pathway, and the RTK-Ras-MAPK pathway (27).

We nucleofected TERT-immortalized NSC-CB660 cells with pools of 3 sgRNAs for each gene, in four successive rounds of nucleofection spaced 1 week apart (to allow cells time to recover from electroporation) (Fig. 3A). In this case, we chose to spread the sgRNAs across each gene to favor individual indels rather than deletions, reasoning that the cumulative effect of 3 sgRNA sites for each gene would lead to a high percentage of cells that contained at least one out-of-frame edit in each allele. We examined the effect on target protein expression via western blotting on the pool at each stage in the process, as well as on eight subclones of the final pool. Remarkably, we observed dramatic protein loss for each gene in the targeted pools (Fig. 3B). Examination of the eight clones of the final *TP53+CDKN2A+PTEN+NF1* targeted KO pool revealed a similar result, where all proteins showed similar reduction in individual clones, except for one clone which still showed protein expression of Nf1 (Fig. 3C).

**Figure 3.**
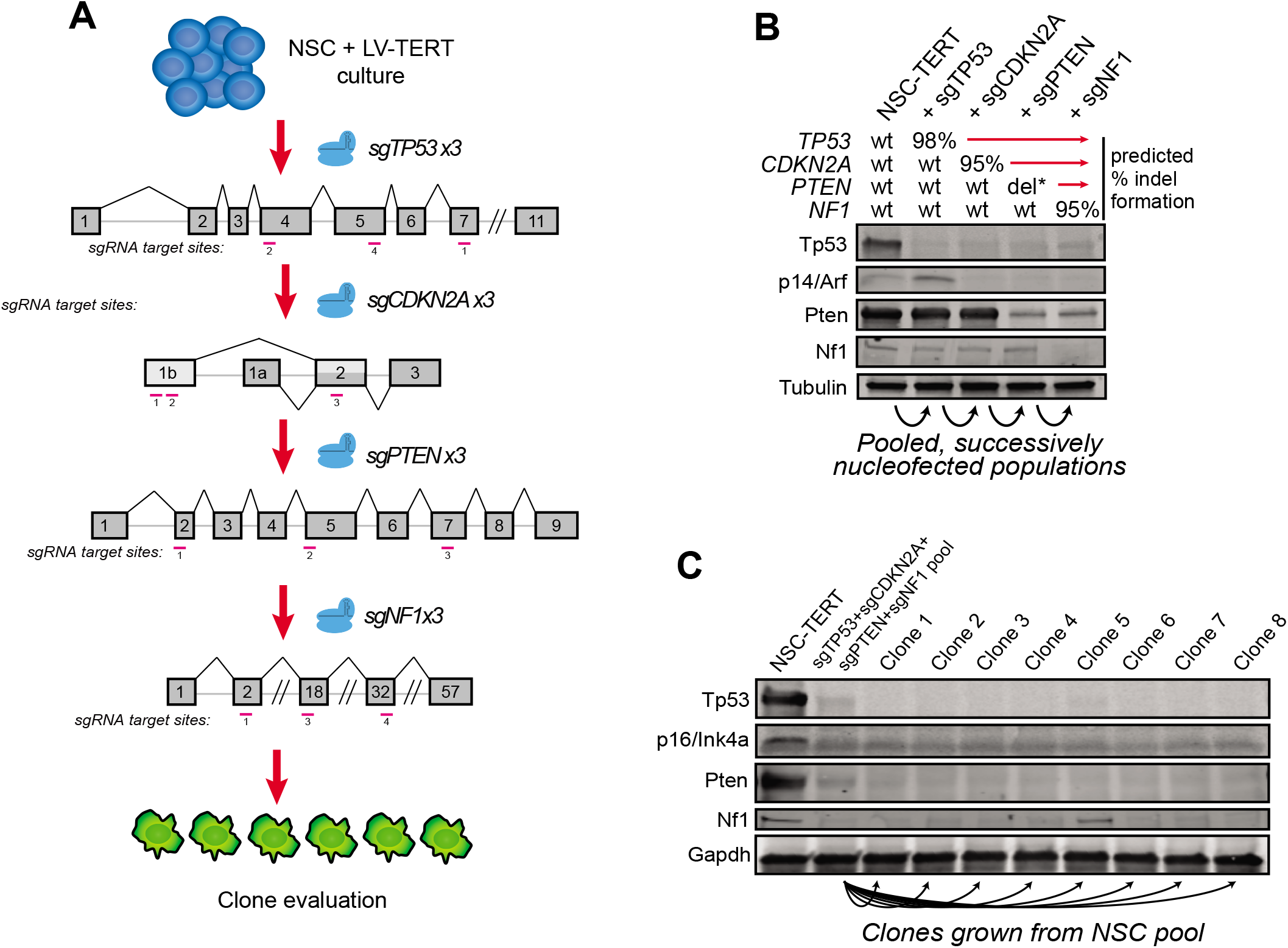
Use of Cas9:sgRNA RNPs to generate multi-gene knockout cell pools to model oncogenic transformation reveals dramatic protein loss. (A) Overview of strategy for successive Cas9:sgRNA RNP nucleofections targeting various tumor suppressors in NSCs. (B) Western blots for targeted genes in the cell pool at each successive stage. Predicted % indel formation for each gene from Sanger sequencing results is shown above blot. del* indicates that large deletions were identified in addition to indels (detailed in Figure 4). For detection of p53, cells were treated with doxorubin to stabilize the protein. (C) Western blots for 8 clones derived from the final sgTP53+sgCDKN2A+ sgPTEN+sgNF1 cell pool.

### RNP nucleofection allows for targeted biallelic deletion of multi-kb genomic regions

We also assessed indel efficiencies in the tumor suppressor KO cell pools across one test sgRNA for each gene. For *TP53, CDKN2A*, and *NF1* we observed high predicted indel frequencies of 98%, 95%, and 95%, respectively. For *PTEN*, however, we noted a discrepancy between the Western blot results, which showed near total ablation of protein expression, and the indel analysis for all 3 sgRNAs, which each revealed only ~33% efficiency. Since probability suggests that even the cumulative editing effect of these 3 sgRNAs should not quite account for near total protein loss, we suspected that the *PTEN* sgRNA pool may have allowed for the generation of a large deletion. To investigate this possibility, we performed PCR with primers flanking the outer sgRNA cut sites (Fig. 4A). In this case, an allele that did *not* harbor deletion of the entire ~64kbp region would not amplify properly in our PCR conditions since the product would be too large. We observed that 2 of 8 clones tested contained a deletion allele, and one of these actually produced two products of similar but distinct sizes, indicating a biallelic deletion with slightly different editing results (Fig. 4B, left panel). As a positive control for gDNA integrity, we used a second PCR spanning a small *PTEN* region outside the deletion region, and we observed the correct product for all 8 clones in this case (Fig. 4B, right panel).

**Figure 4.**
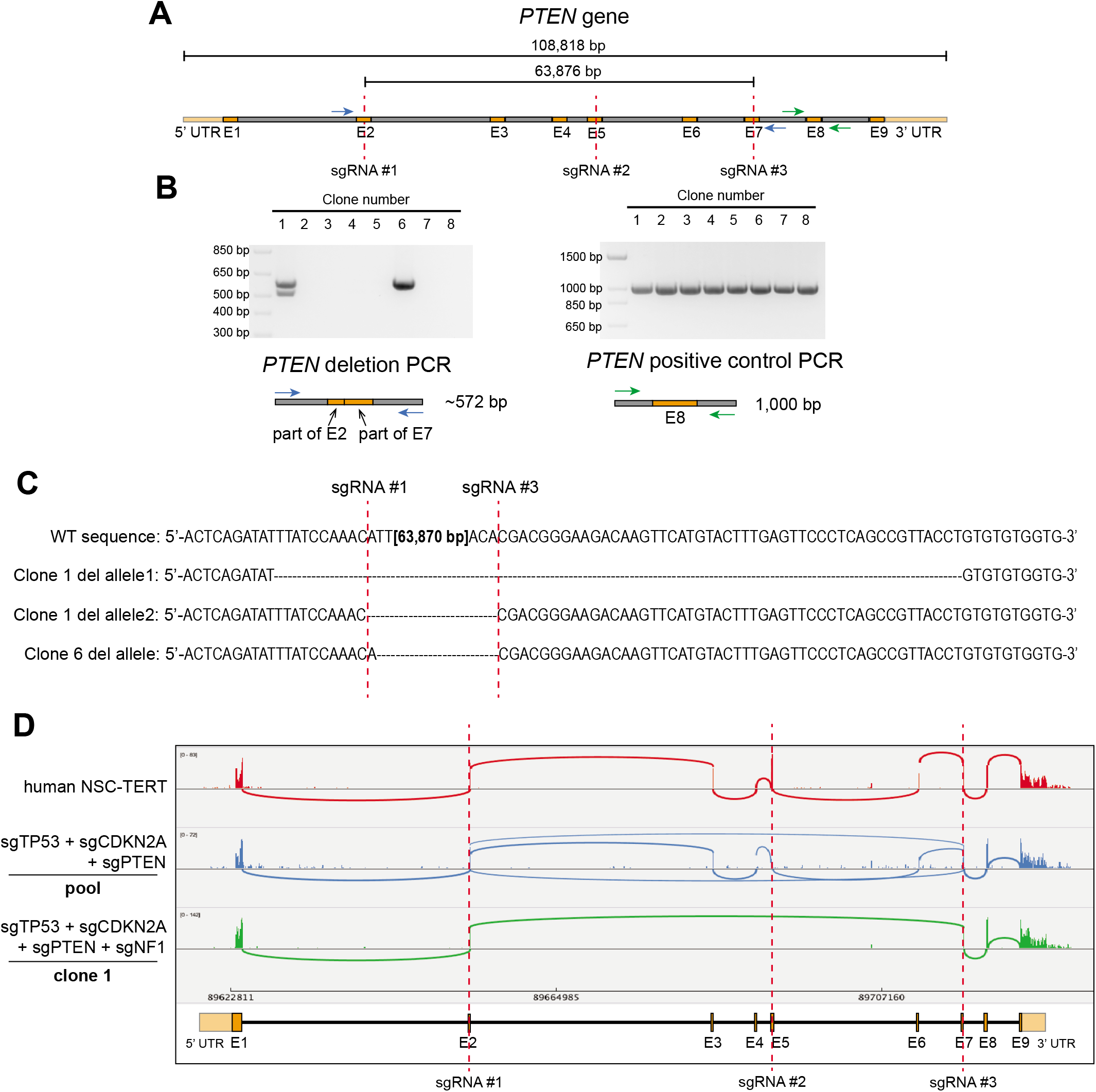
Cas9:sgRNA RNPs can be used to generate large genomic deletions in cell clones. (A) The targeting strategy for the *PTEN* gene. Three sgRNAs were designed targeting exons 2, 5, and 7. To check for the potential deletion of the ~64kb segment between the outermost sgRNAs, deletion PCR primers (in blue) were devised to amplify a product only if the entire region had been deleted. Amplification of the region around the non-targeted exon 8 (primers in green) served as a positive control for gDNA integrity. (B) Left: Deletion PCR as described in (A) for 8 different clones of transformed NSC-CB660 that had been nucleofected with the 3 sgRNAs targeting *PTEN*. Right: Positive control PCR as described in (A) for the 8 clones. (C) Genomic sequences of the 3 deletion alleles identified in (B). Red dotted lines denote sgRNA cut sites. (D) A sashimi plot of RNA-seq reads covering the *PTEN* gene for parental NSC cells, targeted pool, and clone 1. “Transcripts” with a minimum junction coverage of 5 reads are shown.

Gel-purification of the deletion PCR bands followed by Sanger sequencing confirmed that each deletion allele observed in the clones was a result of cutting near the predicted outermost sgRNA cut sites, with one allele tested being an exact deletion, one containing an additional 1bp insertion, and one containing an additional 10bp 5’-deletion and 48bp 3’-deletion (Fig. 4C). To further investigate this, we performed RNA-seq and examined reads mapping to the *PTEN* locus in the KO pool and in clone 1. Analysis of predicted “splice junctions” based on RNA-seq reads showed that mRNA containing the exact deletions observed at the gDNA level could be identified in clone 1, and no properly spliced reads were present, corroborating the fact that this clone did not contain any non-deletion allele (Fig. 4D). Furthermore, reads corresponding to large deletions could be observed in the cell pool as well, in addition to the expected normally spliced reads. These results suggest that using multiple sgRNAs with our method has the potential to create large (>50kbp) deletions, which may be monoallelic or biallelic in subsets of clones.

### Analysis of gene expression changes induced by tumor suppressor KO targeting events

To further assess the fidelity of gene KOs via CRISPR RNP nucleofection, we examined the progressive changes in gene expression after each successive targeting event (sg*TP53*, sg*CDKN2A*, sg*PTEN*, sg*NF1*) in NSC-CB660-TERT cells, by performing RNA-seq on the parental cells compared to the targeted cells at each stage. We used edgeR (28) for differential gene expression analysis, and Figure 5A shows the overall relationship of these data in cluster analysis and the gene expression changes after each targeting. The greatest number of changes were produced by *TP53* KO (269 genes with log2 fold-change (log2FC)>0.5 and 682 genes log2FC<-0.5 (FDR<0.05)) and *CDKN2A* KO (1340 genes log2FC>0.5 and 1733 genes log2FC<-0.5 (FDR<.05)). Importantly, ≥80% of *TP53* KO-induced expression changes were maintained (at FDR<0.05) even after further *CDKN2A* KO, *PTEN* KO, and *NF1* KO, and ≥78% of *CDKN2A* KO induced expression changes were maintained even after further *PTEN* KO and *NF1* KO.

**Figure 5.**
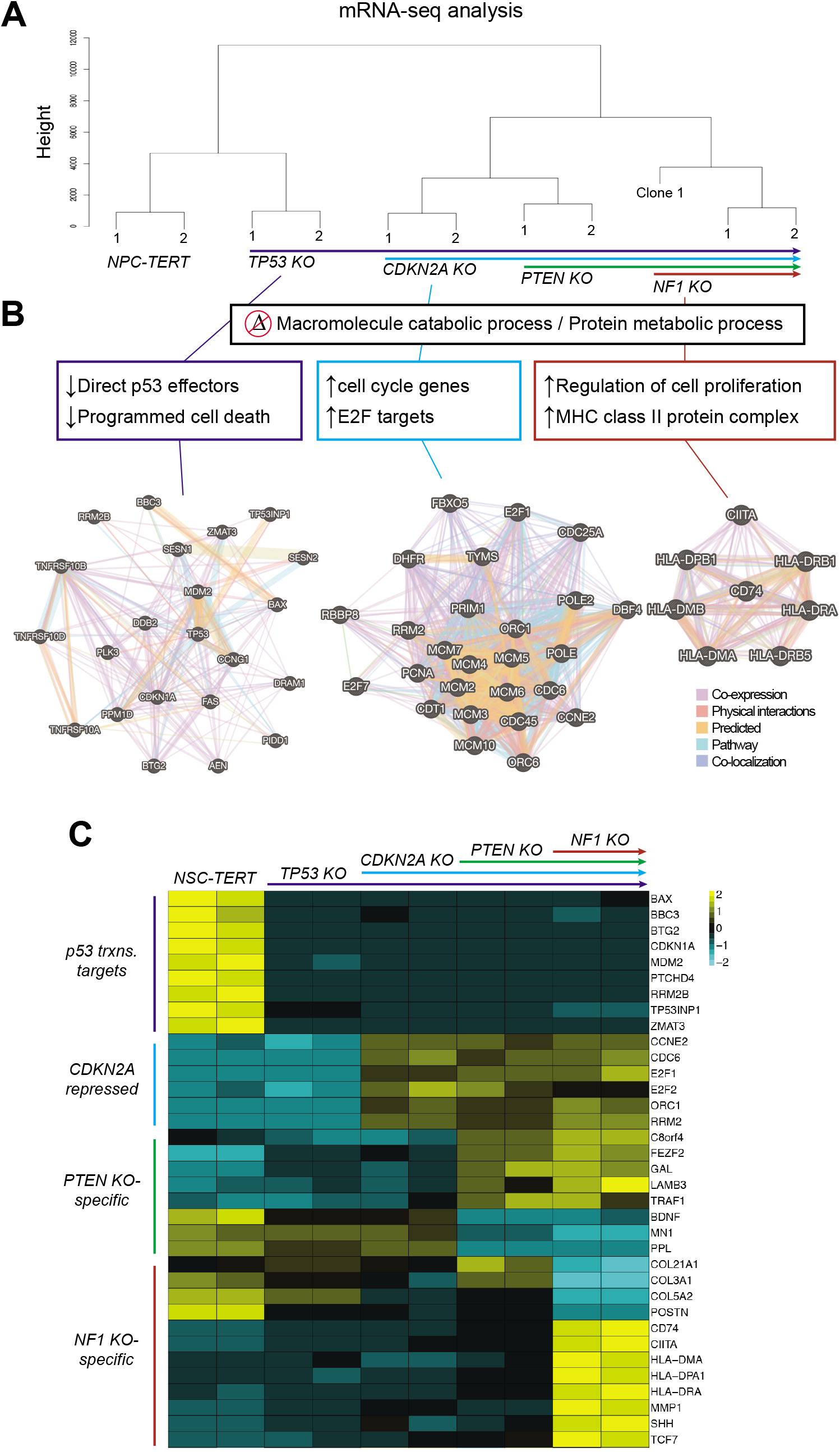
Gene expression changes induced by tumor suppressor knockout targeting events in human NSCs. (A) Cluster analysis showing relative gene expression profile relationships based on RNA-seq for parental NSC-TERT cells and NSC-TERT successively nucleofected with sgTP53, then sgCDKN2A, then sgPTEN, and then sgNF1, as well as for clone 1 derived from the final pool where all 4 genes had been targeted. Duplicates were sequenced for all samples except clone 1. (B) Networks showing gene relationships for genes altered upon TP53, CDKN2A, or NF1 knockout. (C) A heatmap displaying examples of sgTP53, sgCDKN2A, sgPTEN, and sgNF1-specific gene expression changes observed in RNA-seq data. Relative gene expression values were normalized for all samples within each gene. Each RNA-seq replicate is shown as a separate column.

The gene expression changes associated with *TP53* KO cells were consistent with p53’s known transcriptional function. We observed downregulation of key p53 transcriptional targets, including 27 of 132 found in Pathway Interaction Database (p=8.2E-20) and 51 of 116 literature-curated p53 targets, including: *BAX, BBC3/PUMA, BTG2, CDKN1A/p21, RRM2B*, and *ZMAT3* (29) (Fig. 5B & C; Tables S2 & S3). *CDKN2A* KO cells most prominently revealed upregulation of genes associated with the cell cycle (285 genes; GO:0007049; p=6E-116) and specifically *E2F* targets (52 genes; p=2.4E-42), including *E2F1* and *E2F2* themselves (Fig. 5B & C; Tables S2 & S3). This is consistent with loss of *CDKN2A*’s p16 protein, which inhibits Cyclin D/CDK activity in G1, preventing de-repression of E2F (30). Other prominent cell cycle regulated genes included those associated with: *CDK1* interactions (67 genes, p=1.7E-28), *MCM6* interactions (33 genes, p=6.1E-19), *PCNA* interactions (64 genes, p=6.2E-28), and *PLK1* interactions (50 genes, p=6.4E-20). Thus, loss of *CDKN2A* leads to profound reprogramming of the transcription of critical cell cycle-regulated genes in human NSCs, consistent with loss of p16 function.

By contrast to *TP53* and *CDKNA, PTEN* KO produced the fewest changes in gene expression in our scheme (upregulation of 169 genes and downregulation of 216 genes (+/-0.5 log2FC, FDR<0.05)). This may be due to epistasis with gene expression changes caused by *CDKN2A* KO. Nonetheless, manual curation of these genes revealed possible connections to the PI-3 kinase pathway itself, suggestive of feedback regulation. For example, within the downregulated genes, *PPL/Plakin* binds Akt directly (31), and brain-derived neurotrophic factor *(BDNF*) activates Akt (32) and Pten itself (33). Within the upregulated genes, *GAL/galanin* codes for neuroendocrine peptide that exhibits an autocrine mitogenic effect through Erk and Akt activity (34) (Fig. 5C). In addition, there were many novel genes affected by *PTEN* loss, including, among others: *C8orf4/TCIM*, a positive regulator of the Wnt/beta-catenin pathway (34); *FEZF2*, a marker and transcription factor associated with NSC-dependent patterning of the cerebral cortex (35); and *TRAF1*, which mediates the anti-apoptotic signals from TNF receptors (36) (Fig. 5C).

With the further addition of *NF1* KO in *TP53+CDKN2A+PTEN* KO cells, we observed changes in expression of an additional 1022 genes (321 up- and 701 downregulated; log2FC+/- 0.5; FDR<0.05). The downregulated genes were most significantly enriched for extracellular matrix organization genes (43 genes; GO:0030198; p=1.8E-15), which included many collagen encoding genes and Periostin (Fig. 5C). Upregulated genes included genes involved in regulation of cell proliferation (43 genes, p=5.2E-6), including members of the WNT signaling pathway *(WNT7B, FZD4, LEF1*, and *TCF7*). Interestingly, also upregulated were eight genes coding for major histocompatibility complex (MHC) class II proteins, including *CIITA*, the master transcriptional activator controlling expression of the MHC class II genes (37) (Fig. 5B & C; Tables S2 & S3). *CIITA* expression was upregulated ~10-fold in *NF1 KO* cells relative to NSC-CB660-TERT cells. MHC class II genes are primarily expressed by professional antigen presenting cells, such as dendritic cells, macrophages, and B cells (38). However, remarkably, upregulation of MHC class II protein complex is a hallmark of *NF1^-/-^* human neurofibroma tumor cells, which *CIITA* activity is required to maintain (37, 39). It has also been observed that gliomas and other cancers have a high proportion of MHC class II-expressing tumor cells (40), which may promote tolerance to tumor-associated antigens (41). Our results show that *NF1* loss is one route to de-repression of MHC class II machinery in human NSCs.

In addition to gene expression changes in response to loss of *TP53, CDKN2A, PTEN*, and *NF1*, we also identified 806 genes whose expression remained largely unaltered in all conditions (FDR≥0.2 across all comparisons with ≥10 CPM counts across all samples). Gene set enrichment analysis revealed that 487 of these were involved in a “cellular metabolic process” (GO:0044237; FDR=1.5E-15), with 129 encoding mitochondrial proteins (including 19 involved in oxidative phosphorylation) and 29 encoding ribosomal proteins (Tables S2 & S3). Our previous CRISPR-Cas9 lethality screens in NSC-CB660 cells demonstrate that at least 208 of these genes score as essential, including, for example, 34 genes associated with ribosome biogenesis, 11 genes coding for respiratory electron transport machinery, and 23 overlapping with “housekeeping” genes (42) (GSEA: 111197; p= 7.12E-13). These results suggest that a subset of genes expressed in human NSCs are transcriptionally regulated and/or maintain their mRNA levels independently of p53, Rb-axis, PI-3 kinase, and Nf1 pathways.

Altogether, our gene expression results highlight the utility of our CRISPR/Cas9 RNP nucleofection method for quickly creating a series of KOs that allow for the study of gene activities. Importantly, due to the high targeting efficiency using our method, we were able to confirm many known as well as identify novel transcriptional changes associated with loss of the genes we targeted, using nucleofected cell *pools* rather than cell clones.

## METHODS

### CRISPR sgRNA Design

CRISPR sgRNAs were designed via manual curation of all possible sgRNA sequences for a given region as identified by the Broad Institute’s GPP Web Portal (26). See Table S1 for a list of sgRNA sequences used.

### Cas9:sgRNA RNP Nucleofection

See detailed protocol in Supplementary Materials. Briefly, to prepare RNP complexes, reconstituted sgRNA (Synthego) and then sNLS-SpCas9-sNLS (Aldevron) were added to complete SG Cell Line Nucleofector Solution (Lonza), to a final volume of 20 μL. The mixture was incubated at room temperature for 15 minutes to allow RNP complexes to form. A Cas9:sgRNA molar ratio of 1:2 was used, unless otherwise noted. Total RNP doses described refer to the amount of the limiting complex member (Cas9). To nucleofect, 1.5 x 10^5^ cells were harvested, washed with PBS, resuspended in 20 μL of RNPs, and electroporated using the Amaxa 96-well Shuttle System or 4D X Unit (Lonza) and program EN-138.

### CRISPR Editing Analysis

Nucleofected cells were harvested at indicated timepoints and genomic DNA was extracted (MicroElute Genomic DNA Kit, Omega Bio-Tek). Genomic regions around CRISPR target sites were PCR amplified using Phusion polymerase (Thermo Fisher) and primers located (whenever possible) at least 250bp outside cut sites. After size verification by agaorse gel electrophoresis, PCR products were column-purified (Monarch PCR & DNA Clean-up Kit, New England BioLabs) and submitted for Sanger sequencing (Genewiz) using unique sequencing primers. The resulting trace files for edited cells versus control cells (nucleofected with non-targeting Cas9:sgRNA) were analyzed for predicted indel composition using the Inference of CRISPR Edits (ICE) web tool (25). See Table S1 for a list of all PCR and sequencing primers used.

### Cell Culture

Patient tumor-derived GSCs and fetal tissue-derived NSCs were provided by Drs. Do-Hyun Nam, Jeongwu Lee, and Steven M. Pollard, were obtained via informed consent, and have been previously published (49-51). Isolates were cultured in NeuroCult NS-A basal medium (StemCell Technologies) supplemented with B27 (Thermo Fisher), N2 (homemade 2x stock in Advanced DMEM/F-12 (Thermo Fisher)), EGF and FGF-2 (20 ng/ml) (PeproTech), glutamax (Thermo Fisher), and antibiotic-antimycotic (Thermo Fisher). Cells were cultured on laminin (Trevigen or in-house-purified) -coated polystyrene plates and passaged as previously described (49), using Accutase (EMD Millipore) to detach cells.

### Western blotting

Cells were harvested, washed with PBS, and lysed with modified RIPA buffer (150mM NaCl, 25mM Tris-HCl (pH 8.0), 1mM EDTA, 1.0% Igepal CA-630 (NP-40), 0.5% sodium deoxycholate, 0.1% SDS, 1X protease inhibitor cocktail (complete Mini EDTA-free, Roche)). Lysates were sonicated (Bioruptor, Diagenode) and then quantified using Pierce BCA assay (Thermo Fisher). Identical amounts of proteins (20-40μg) were electrophoresed on 4–15% Mini-PROTEAN TGX precast protein gels (Bio-Rad). For transfer, the Trans-Blot Turbo transfer system (Bio-Rad) with nitrocellulose membranes was used according to the manufacturer’s instructions. TBS (137mM NaCl, 20mM Tris, pH 7.6) +5% nonfat milk was used for blocking, and TBS+0.1%Tween-20+5% milk was used for antibody incubations. The following commercial primary antibodies were used: Tp53 (Cell Signaling #48818, 1:500), p14/Arf (Bethyl Laboratories #A300-340A-T, 1:500), p16/Ink4a (Abcam #ab16123, 1:200), Pten (Cell Signaling #9559S, 1:1,000), Nf1 (Santa Cruz #sc-67, 1:50), aTubulin (Sigma #T9026, 1:1,000), Gapdh (Sigma #SAB2500450, 1:100). The following secondary antibodies were used (LI-COR): #926-68073, #926-32212, #926-32214, #926-68074. An Odyssey infrared imaging system (LI-COR) was used to visualize blots.

### RNA-seq and analysis

Cells were lysed with Trizol (Thermo Fisher). Total RNA was isolated (Direct-zol RNA kit, Zymo Research) and quality validated on the Agilent 2200 TapeStation. Illumina sequencing libraries were generated with the KAPA Biosystems Stranded RNA-Seq Kit (52) and sequenced using HiSeq 2000 (Illumina) with 100bp paired-end reads. RNA-seq reads were aligned to the UCSC hg19 assembly using STAR2 (v 2.6.1) (53) and counted for gene associations against the UCSC genes database with HTSeq (54). Normalized gene count data was used for subsequent hierarchical clustering (R package ggplot2 (55)) and differential gene expression analysis (R/Bioconductor package edgeR (28)). Heatmaps were made using R package pheatmap (56).

## DISCUSSION

Here, we present a method for creating bi- and multi-allelic loss-of-function indel mutations, using *in vitro* assembled Cas9:sgRNA RNPs composed of chemically synthesized, modified sgRNA and purified Cas9 protein. By this method, indel efficiencies of >90% can routinely be achieved in populations of cells, obviating the need for clonal selection of edited cells or chemical selection of DNA-based sgRNA expression systems. Moreover, because gene editing is complete within three days of RNP introduction, this approach offers better experimental tractability over current approaches, which can suffer from lack of indel penetrance and protracted windows of indel formation.

This method also improves upon existing methods for the creation of precise or near-precise deletions. In mammals, single dsDNA breaks, including those generated by Cas9:sgRNA cutting, produce NHEJ-dependent small insertions or deletions at break sites (i.e., error-prone repair) (11, 43). However, adding a second dsDNA break in close proximity to the first can cause rejoining without error via “accurate” NHEJ (44, 45). Our results support this notion. We observe a high frequency and penetrance of conversion of single indels to precise and near-precise deletions. It has been previously shown that using 2-3 proximal sgRNAs can create deletions of ~10bp to 1Mb in mouse embryos and cultured cells (4, 44, 46, 47). However, these approaches produce deletion formation efficiencies ranging from ~2% to ~40%. By contrast, we observe near-precise deletion frequencies as high as >90% using sgRNAs spaced up to 1000bp apart. This suggests that the use of 2 sgRNAs using our approach can have the added benefit of triggering accurate NHEJ and being able to specify a high frequency of precise or near-precise deletions.

This technique does have a few limitations. First, achieving high multi-allelic indel efficiencies may require pre-validation of sgRNAs. However, we have had good success designing sgRNAs using the Broad GPP Web Portal design tool (where ~60-70% of sgRNAs that we choose via manual curation produce >80% indel formation) or, alternatively, choosing sgRNA sequences from positively scoring CRISPR-Cas9 lentiviral-based screen hits. Second, reliance on chemically synthesized sgRNAs can be cost-prohibitive for large-scale projects. An alternative option is to generate *in vitro* transcribed sgRNA using T7, T3, or SP6 RNA polymerase in the presence of ribonucleoside triphosphates and a DNA template (3). However, this requires additional steps, namely the initial creation of an accurate DNA template and the purification of the transcribed sgRNA to remove unincorporated triphosphates, enzyme, and template DNA. In addition, *in vitro* transcription can result in errors toward the 5’ end of the RNA (48). Also, it is not possible to easily generate modifications, meaning *in vitro* transcribed sgRNA does not possess increased protection from nucleases once it has entered the cell, resulting in decreased editing efficiency compared to chemically synthesized, modified sgRNA. Nonetheless, it is a viable alternative to consider when cost is a concern. Another alternative is to employ a commercially sourced dual gRNA system (crRNA:tracrRNA), which may represent only a slight reduction in efficiency. The two chemically synthesized RNAs can still be modified to enhance nuclease resistance, but they are usually available at a lower cost since the accurate synthesis of these shorter RNAs is less complex compared to a longer chimeric sgRNA.

Our data suggest that the relatively simple method described here can be used for highly efficient (>90%) and fast (72 hours) gene knockout, as well as for targeted genomic deletions, even in hyperdiploid cells (such as many tumor cells). This represents an extremely useful tool for inactivating not only coding genes, but also non-coding elements such as non-coding RNAs, UTRs, enhancers, and promoters. The gain in efficiency that we observe can allow for systematic well-by-well screening (similar to small interfering RNA screens), but provides the flexibility of targeting any small (<1000bp) element in the genome. This method can be readily applied to diverse mammalian cell types by varying the nucleofection buffer and program (Lonza can provide appropriate conditions for many cell types). Thus, it represents an important step forward in the ability to manipulate the genomes of cell cultures derived from primary cells, such as patient-derived tumor cells and human stem/progenitor cells.

## Supporting information

Supplemental Protocol

## ACKNOWLEDGEMENTS

We thank members of the Paddison laboratory for helpful discussions and critical reading of the manuscript; Do-Hyun Nam, Jeongwu Lee, and Steven M. Pollard for providing cell isolates; Franziska Michor for helpful suggestions; David McDonald in the Cellular Imaging Core for performing karyotyping analysis; and Pam Lindberg and An Tyrrell for administrative support. *Funding Sources:* National Institutes of Health (T32 CA080416 to P.H., R01 CA190957, P30 DK56465-13, U54 DK106829, P30 CA015704); American Cancer Society (ACS-RSG-14-056-01); Robert J. Kleberg, Jr. and Helen C. Kleberg Foundation.

## AUTHORS’ CONTRIBUTIONS

PH, MK, H-JW, HMF, and PJP designed the experiments. PH and MK performed the majority of experiments and analyzed the corresponding data. SA performed bioinformatics processing and analysis of RNA-seq data. HMF assisted with experiments. PH and PJP wrote the manuscript. All authors read and approved the final manuscript.

## DATA AVAILABILITY STATEMENT

The data that supports the findings of this study are available in the supplementary material of this article.

## REFERENCES

1. Wiedenheft B, Sternberg SH, Doudna JA. RNA-guided genetic silencing systems in bacteria and archaea. Nature. 2012;482(7385):331–8.

2. Mali P, Esvelt KM, Church GM. Cas9 as a versatile tool for engineering biology. Nat Methods. 2013;10(10):957–63.

3. Cho SW, Kim S, Kim JM, Kim JS. Targeted genome engineering in human cells with the Cas9 RNA-guided endonuclease. Nat Biotechnol. 2013;31(3):230–2.

4. Cong L, Ran FA, Cox D, Lin S, Barretto R, Habib N, et al. Multiplex Genome Engineering Using CRISPR/Cas Systems. Science. 2013;339:819–23.

5. Jinek M, East A, Cheng A, Lin S, Ma E, Doudna J. RNA-programmed genome editing in human cells. Elife. 2013;2:e00471.

6. Mali P, Yang L, Esvelt KM, Aach J, Guell M, DiCarlo JE, et al. RNA-guided human genome engineering via Cas9. Science. 2013;339:823–6.

7. Jinek M, Chylinski K, Fonfara I, Hauer M, Doudna JA, Charpentier E. A Programmable Dual-RNA-Guided DNA Endonuclease in Adaptive Bacterial Immunity. Science. 2012;337:816–21.

8. Ran FA, Hsu PD, Wright J, Agarwala V, Scott DA, Zhang F. Genome engineering using the CRISPR-Cas9 system. Nature Protocols. 2013;8:2281–308.

9. Hartlerode AJ, Scully R. Mechanisms of double-strand break repair in somatic mammalian cells. The Biochemical journal. 2009;423(2):157–68.

10. Lieber MR, Ma Y, Pannicke U, Schwarz K. Mechanism and regulation of human non-homologous DNA end-joining. Nat Rev Mol Cell Biol. 2003;4(9):712–20.

11. Chakrabarti AM, Henser-Brownhill T, Monserrat J, Poetsch AR, Luscombe NM, Scaffidi P. Target-Specific Precision of CRISPR-Mediated Genome Editing. Mol Cell. 2019;73(4):699–713 e6.

12. Shalem O, Sanjana NE, Hartenian E, Shi X, Scott DA, Mikkelsen TS, et al. Genomescale CRISPR-Cas9 knockout screening in human cells. Science. 2014;343:84–7.

13. Wang T, Wei JJ, Sabatini DM, Lander ES. Genetic screens in human cells using the CRISPR-Cas9 system. Science. 2014;343:80–4.

14. Toledo CM, Ding Y, Hoellerbauer P, Davis RJ, Basom R, Girard EJ, et al. Genome-wide CRISPR-Cas9 Screens Reveal Loss of Redundancy between PKMYT1 and WEE1 in Glioblastoma Stem-like Cells. Cell Rep. 2015;13(11):2425–39.

15. Kim S, Kim D, Cho SW, Kim J, Kim JS. Highly efficient RNA-guided genome editing in human cells via delivery of purified Cas9 ribonucleoproteins. Genome Research. 2014;24(6):1012–9.

16. Zuris JA, Thompson DB, Shu Y, Guilinger JP, Bessen JL, Hu JH, et al. Cationic lipid-mediated delivery of proteins enables efficient protein-based genome editing in vitro and in vivo. Nature Biotechnology. 2014;33:73.

17. DeWitt MA, Corn JE, Carroll D. Genome editing via delivery of Cas9 ribonucleoprotein. Methods. 2017;121-122:9–15.

18. Schumann K, Lin S, Boyer E, Simeonov DR, Subramaniam M, Gate RE, et al. Generation of knock-in primary human T cells using Cas9 ribonucleoproteins. Proceedings of the National Academy of Sciences. 2015;112(33):10437.

19. Lin S, Staahl BT, Alla RK, Doudna JA. Enhanced homology-directed human genome engineering by controlled timing of CRISPR/Cas9 delivery. eLife. 2014;3:e04766.

20. Bressan RB, Dewari PS, Kalantzaki M, Gangoso E, Matjusaitis M, Garcia-Diaz C, et al. Efficient CRISPR/Cas9-assisted gene targeting enables rapid and precise genetic manipulation of mammalian neural stem cells. Development. 2017;144:635–48.

21. Graf R, Li X, Chu VT, Rajewsky K. sgRNA Sequence Motifs Blocking Efficient CRISPR/Cas9-Mediated Gene Editing. Cell Reports. 2019;26(5):1098–103.e3.

22. Eckstein F. Phosphorothioates, Essential Components of Therapeutic Oligonucleotides. Nucleic Acid Therapeutics. 2014;24(6):374–87.

23. Allerson CR, Sioufi N, Jarres R, Prakash TP, Naik N, Berdeja A, et al. Fully 2’-Modified Oligonucleotide Duplexes with Improved in Vitro Potency and Stability Compared to Unmodified Small Interfering RNA. Journal of Medicinal Chemistry. 2005;48(4):901–4.

24. Brinkman EK, Chen T, Amendola M, van Steensel B. Easy quantitative assessment of genome editing by sequence trace decomposition. Nucleic Acids Res. 2014;42(22):e168.

25. Hsiau T, Conant D, Maures T, Waite K, Yang J, Kelso R, et al. Inference of CRISPR Edits from Sanger Trace Data. bioRxiv. 2019.

26. Doench JG, Fusi N, Sullender M, Hegde M, Vaimberg EW, Donovan KF, et al. Optimized sgRNA design to maximize activity and minimize off-target effects of CRISPR-Cas9. Nat Biotechnol. 2016;34(2):184–91.

27. Brennan CW, Verhaak RGW, McKenna A, Campos B, Noushmehr H, Salama SR, et al. The somatic genomic landscape of glioblastoma. Cell. 2013;155:462–77.

28. Robinson MD, McCarthy DJ, Smyth GK. edgeR: a Bioconductor package for differential expression analysis of digital gene expression data. Bioinformatics (Oxford, England). 2010;26(1):139–40.

29. Fischer M. Census and evaluation of p53 target genes. Oncogene. 2017;36(28):3943–56.

30. Sherr CJ, McCormick F. The RB and p53 pathways in cancer. Cancer Cell. 2002;2(2):103–12.

31. van den Heuvel AP, de Vries-Smits AM, van Weeren PC, Dijkers PF, de Bruyn KM, Riedl JA, et al. Binding of protein kinase B to the plakin family member periplakin. J Cell Sci. 2002;115(Pt 20):3957–66.

32. Song W, Volosin M, Cragnolini AB, Hempstead BL, Friedman WJ. ProNGF induces PTEN via p75NTR to suppress Trk-mediated survival signaling in brain neurons. J Neurosci. 2010;30(46):15608–15.

33. Plotkin JL, Day M, Peterson JD, Xie Z, Kress GJ, Rafalovich I, et al. Impaired TrkB receptor signaling underlies corticostriatal dysfunction in Huntington’s disease. Neuron. 2014;83(1):178–88.

34. Banerjee R, Henson BS, Russo N, Tsodikov A, D’Silva NJ. Rap1 mediates galanin receptor 2-induced proliferation and survival in squamous cell carcinoma. Cell Signal. 2011;23(7):1110–8.

35. Dwyer ND, Chen B, Chou SJ, Hippenmeyer S, Nguyen L, Ghashghaei HT. Neural Stem Cells to Cerebral Cortex: Emerging Mechanisms Regulating Progenitor Behavior and Productivity. J Neurosci. 2016;36(45):11394–401.

36. Shu HB, Takeuchi M, Goeddel DV. The tumor necrosis factor receptor 2 signal transducers TRAF2 and c-IAP1 are components of the tumor necrosis factor receptor 1 signaling complex. Proceedings of the National Academy of Sciences of the United States of America. 1996;93(24):13973–8.

37. Choi NM, Majumder P, Boss JM. Regulation of major histocompatibility complex class II genes. Curr Opin Immunol. 2011;23(1):81–7.

38. Neefjes J, Jongsma ML, Paul P, Bakke O. Towards a systems understanding of MHC class I and MHC class II antigen presentation. Nat Rev Immunol. 2011;11(12):823–36.

39. Reuss DE, Mucha J, Holtkamp N, Muller U, Berlien HP, Mautner VF, et al. Functional MHC class II is upregulated in neurofibromin-deficient Schwann cells. J Invest Dermatol. 2013;133(5):1372–5.

40. Tran CT, Wolz P, Egensperger R, Kosel S, Imai Y, Bise K, et al. Differential expression of MHC class II molecules by microglia and neoplastic astroglia: relevance for the escape of astrocytoma cells from immune surveillance. Neuropathol Appl Neurobiol. 1998;24(4):293–301.

41. Thibodeau J, Bourgeois-Daigneault MC, Lapointe R. Targeting the MHC Class II antigen presentation pathway in cancer immunotherapy. Oncoimmunology. 2012;1(6):908–16.

42. Hsiao LL, Dangond F, Yoshida T, Hong R, Jensen RV, Misra J, et al. A compendium of gene expression in normal human tissues. Physiol Genomics. 2001;7(2):97–104.

43. Symington LS, Gautier J. Double-strand break end resection and repair pathway choice. Annu Rev Genet. 2011;45:247–71.

44. Guo T, Feng YL, Xiao JJ, Liu Q, Sun XN, Xiang JF, et al. Harnessing accurate non-homologous end joining for efficient precise deletion in CRISPR/Cas9-mediated genome editing. Genome Biol. 2018;19(1):170.

45. Betermier M, Bertrand P, Lopez BS. Is non-homologous end-joining really an inherently error-prone process? PLoS Genet. 2014;10(1):e1004086.

46. Canver MC, Bauer DE, Dass A, Yien YY, Chung J, Masuda T, et al. Characterization of genomic deletion efficiency mediated by clustered regularly interspaced short palindromic repeats (CRISPR)/Cas9 nuclease system in mammalian cells. J Biol Chem. 2014;289(31):21312–24.

47. Wolfs JM, Hamilton TA, Lant JT, Laforet M, Zhang J, Salemi LM, et al. Biasing genomeediting events toward precise length deletions with an RNA-guided TevCas9 dual nuclease. Proceedings of the National Academy of Sciences of the United States of America. 2016;113(52):14988–93.

48. Helm M, Brulé H, Giegé R, Florentz C. More mistakes by T7 RNA polymerase at the 5’ ends of in vitro-transcribed RNAs. RNA. 1999;5:618–21.

49. Pollard SM, Yoshikawa K, Clarke ID, Danovi D, Stricker S, Russell R, et al. Glioma stem cell lines expanded in adherent culture have tumor-specific phenotypes and are suitable for chemical and genetic screens. Cell Stem Cell. 2009;4:568–80.

50. Sun Y, Pollard S, Conti L, Toselli M, Biella G, Parkin G, et al. Long-term tripotent differentiation capacity of human neural stem (NS) cells in adherent culture. Molecular and cellular neurosciences. 2008;38:245–58.

51. Lee J, Kotliarova S, Kotliarov Y, Li A, Su Q, Donin NM, et al. Tumor stem cells derived from glioblastomas cultured in bFGF and EGF more closely mirror the phenotype and genotype of primary tumors than do serum-cultured cell lines. Cancer Cell. 2006;9:391–403.

52. Hart T, Chandrashekhar M, Aregger M, Durocher D, Angers S, Moffat J, et al. High-Resolution CRISPR Screens Reveal Fitness Genes and Genotype-Specific Cancer Liabilities Screens Reveal Fitness Genes. Cell. 2015:1–12.

53. Dobin A, Davis CA, Schlesinger F, Drenkow J, Zaleski C, Jha S, et al. STAR: ultrafast universal RNA-seq aligner. Bioinformatics (Oxford, England). 2013;29(1):15–21.

54. Anders S, Pyl PT, Huber W. HTSeq--a Python framework to work with high-throughput sequencing data. Bioinformatics (Oxford, England). 2015;31(2):166–9.

55. Wickham H, Sievert C, Springer International Publishing AG. ggplot2: elegant graphics for data analysis 2016.

56. Kolde R. pheatmap: Pretty Heatmaps. R package version 1.0.10. https://CRANR-projectorg/package=pheatmap. 2018.

