## Supplemental Figures for "A simple and highly efficient method for multi-allelic CRISPR-Cas9 editing in primary cell cultures"

***Inventory of Supporting Information***

**Supplementary Figures…………………………………………………………………………...……2**

Figure S1: Additional sgRNA targeting for different loci and different cell isolates.

Figure S2: Further evidence that CRISPR RNPs can be used to create near-precise targeted deletions.

Figure S3: CRISPR RNPs allow for targeted deletion of up to ~1000bp genomic regions.

**Detailed protocol: *Cas9:sgRNA ribonucleoprotein nucleofection*…….…….see separate file**

**Supplementary Tables………………………………………………………….see separate files**

Table S1: sgRNA sequences, PCR primer sequences, and PCR conditions

Table S2: RNAseq edgeR analysis_CB660TERT comparison

Table S3: RNAseq edgeR analysis_gene to gene comparison

### Figure S1

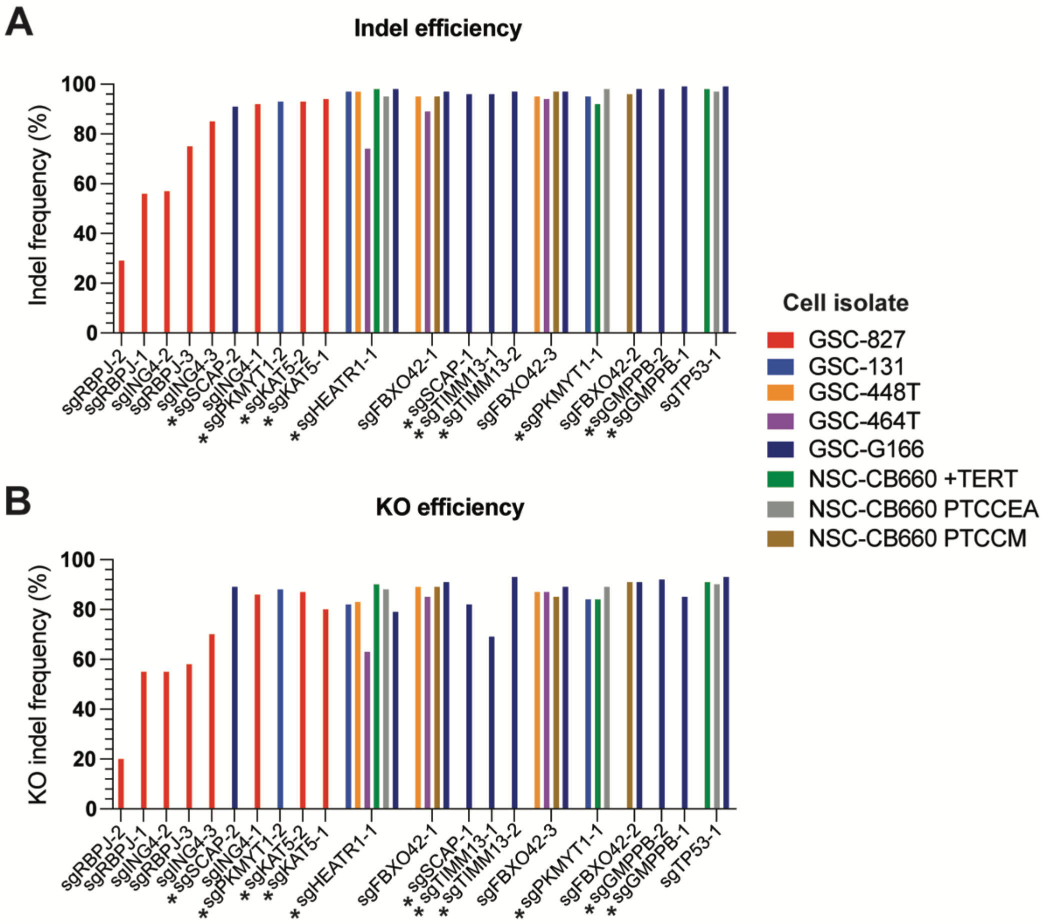


**Figure S1. Additional sgRNA targeting for different loci and different cell isolates.** (A) Predicted indel frequencies for 21 different sgRNAs tested in various GSC and NSC cell isolates. Gene knockout using sgRNAs labeled with * caused reduced viability and/or reduced proliferation in the cell isolate(s) shown. +TERT means cells were transduced with a retroviral vector expressing TERT. +PTCCEA means cells were transduced with dominant-negative TP53DD, CCND1 + CDK4R24C (p16 binding deficient), EGFRvIII, and myristoylated AKT. +PTCCM means cells were transduced with dominant-negative TP53DD, CCND1 + CDK4R24C (p16 binding deficient), and c-Myc. (B) Corresponding predicted knockout frequencies for the sgRNAs shown in (A).

#### Figure S2

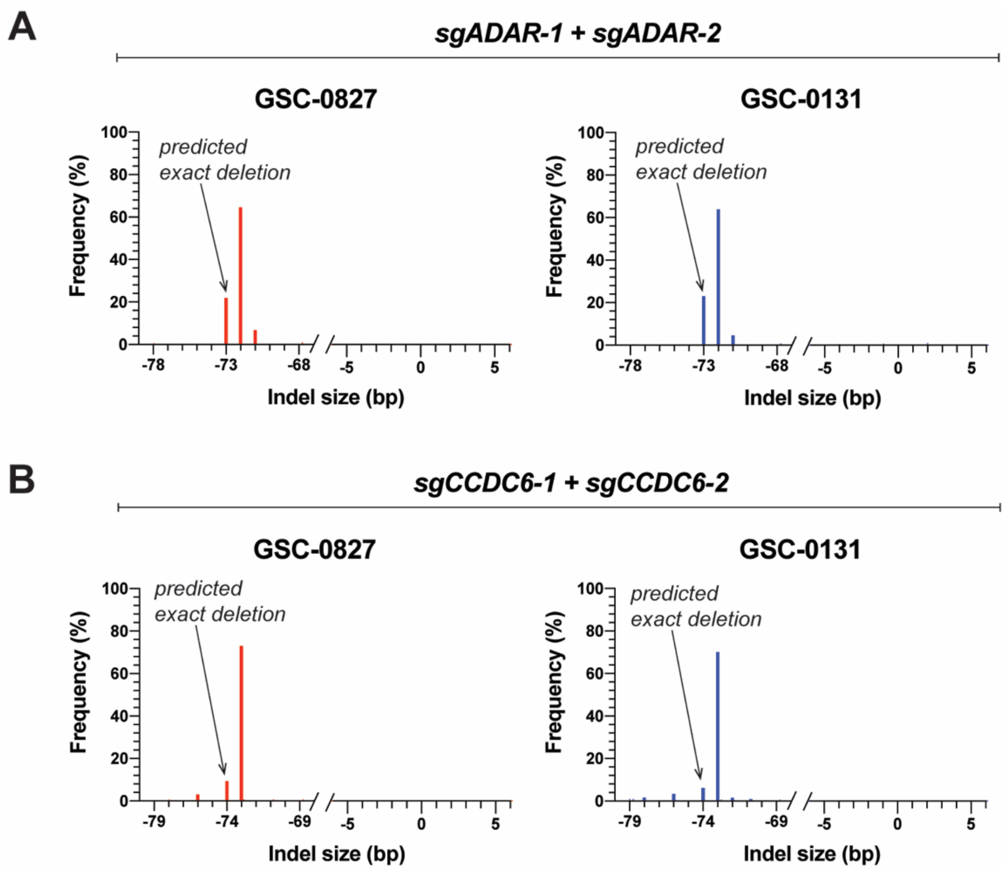


**Figure S2. Further evidence that CRISPR RNPs can be used to create near-precise targeted deletions.** (A) Indel size distribution of predicted indel sequences for 2 GSC cell isolates nucleofected simultaneously with two sgRNAs targeting ADAR. The dual sgRNA cut sites were 73 bp apart. (B) Indel size distribution of predicted indel sequences for 2 GSC cell isolates nucleofected simultaneously with two sgRNAs targeting CCDC6. The dual sgRNA cut sites were 74 bp apart.

### Figure S3

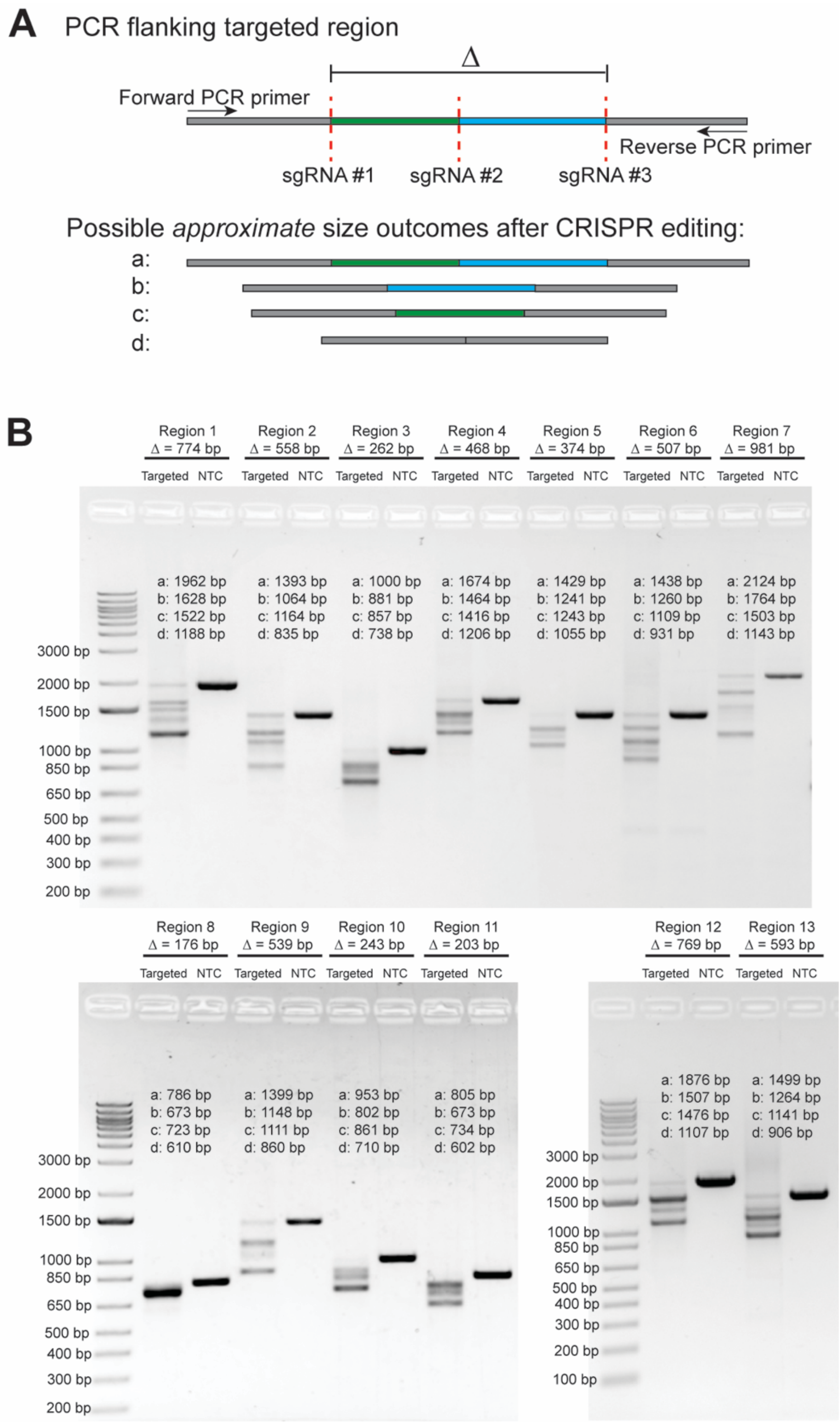
A
PCR flanking targeted region
Δ
Forward PCR primer
Reverse PCR primer
sgRNA #1
sgRNA #2
sgRNA #3
Possible approximate size outcomes after CRISPR editing:
a:
b:
c:
d:
B
Region 1 Δ = 774 bp
Region 2 Δ = 558 bp
Region 3 Δ = 262 bp
Region 4 Δ = 468 bp
Region 5 Δ = 374 bp
Region 6 Δ = 507 bp
Region 7 Δ = 981 bp
Targeted NTC
a: 1962 bp b: 1628 bp c: 1522 bp d: 1188 bp
a: 1393 bp b: 1064 bp c: 1164 bp d: 835 bp
a: 1000 bp b: 881 bp c: 857 bp d: 738 bp
a: 1674 bp b: 1464 bp c: 1416 bp d: 1206 bp
a: 1429 bp b: 1241 bp c: 1243 bp d: 1055 bp
a: 1438 bp b: 1260 bp c: 1109 bp d: 931 bp
a: 2124 bp b: 1764 bp c: 1503 bp d: 1143 bp
3000 bp
2000 bp
1500 bp
1000 bp
850 bp
650 bp
500 bp
400 bp
300 bp
200 bp
Region 8 Δ = 176 bp
Region 9 Δ = 539 bp
Region 10 Δ = 243 bp
Region 11 Δ = 203 bp
a: 786 bp b: 673 bp c: 723 bp d: 610 bp
a: 1399 bp b: 1148 bp c: 1111 bp d: 860 bp
a: 953 bp b: 802 bp c: 861 bp d: 710 bp
a: 805 bp b: 673 bp c: 734 bp d: 602 bp
Region 12 Δ = 769 bp
Region 13 Δ = 593 bp
a: 1876 bp b: 1507 bp c: 1476 bp d: 1107 bp
a: 1499 bp b: 1264 bp c: 1141 bp d: 906 bp
100 bp

**Figure S3. CRISPR RNPs allow for targeted deletion of up to ~1000bp genomic regions.** (A) Schematic showing targeting strategy for deletion of various genomic regions (not to scale). Three sgRNAs were designed for each region, and sgRNA cut sites are denoted by red dotted lines. Δ denotes largest deletion, between outermost sgRNAs. As shown, PCR amplicons of varying sizes are predicted to result from deletion of the region between all 3 sgRNAs (outcome "d"), deletions between only two of the 3 sgRNAs (outcomes "b" and "c"), and no large deletion (wt allele or only small indels; outcome "a"). (B) PCRs for scheme shown in (A) for 13 different genomic regions, which were each targeted in GSC-0827 cells using a total of 60 pmoles RNPs, with a Cas9:sgRNA ratio of 1:3. Cells were harvested for gDNA extraction 5 days post nucleofection. Each region's PCR is shown for a targeted sample compared to a sample nucleofected with a non-targeting control sgRNA RNP (60 pmoles). Predicted sizes of the various PCR amplicons a-d described in (A), as well as the predicted size of the outermost deletion Δ, are noted for each region.
