## Supplemental Protocol for "A simple and highly efficient method for multi-allelic CRISPR-Cas9 editing in primary cell cultures"

### <u>Protocol for Cas9:sgRNA ribonucleoprotein nucleofection using Amaxa 4D Nucleofector X Unit or 96-well Shuttle System</u>

The goal of this protocol is to produce bi- or multi-allelic loss of gene function in cell populations via insertion-deletion (indel) or near-precise deletion mutations triggered by CRISPR-Cas9 targeting. This protocol represents our current preferred method, which utilizes *in vitro*-formed Cas9:sgRNA ribonucleoprotein (RNP) complexes composed of one to three chemically synthesized 2'-O-methyl 3'phosphorothioate-modified sgRNAs and purified Cas9 protein. We currently purchase our sgRNAs from Synthego (https://www.synthego.com/). These sgRNAs are a single RNA molecule that contains both the custom-designed short crRNA sequence fused to the scaffold tracrRNA sequence. Synthego's 2'-O-methyl 3'phosphorothioate-modifications occur in the first and last 3 nucleotides. Other companies also provide sgRNA synthesis services, including IDT, Dharmacon, Thermo Fisher Scientific, and Sigma-Aldrich. In addition, sgRNAs can be synthesized *in vitro*, e.g. using *in vitro* transcription reactions and short DNA templates (although this usually results in inferior editing efficiency).

The Amaxa Nucleofector System uses proprietary electrical parameters and buffer formulations. This protocol is optimized for glioblastoma stem-like cells (GSCs) and human neural stem cells (NSCs) (with nucleofection program based on conditions described for mouse NSCs in Bressan et al. 2017). However, we and collaborators have successfully employed CRISPR in other cell types as well using this method. *Nucleofection conditions (i.e. nucleofection buffer and program) should be optimized for other cell types.* Lonza can provide guidance for many cell lines/types (https://knowledge.lonza.com/).

1. <u>Designing sgRNAs:</u>

   a. There are many sgRNA design resources available. We typically use the Broad Institute's GPP Web Portal (Doench et al. 2016, https://portals.broadinstitute.org/gpp/public/analysis-tools/sgrna-design) in combination with Synthego's design tool (https://design.synthego.com/#/) to identify potential top sgRNAs for a particular gene. We then manually curate these lists by evaluating on-target score versus predicted off-target effects in order to identify ~3-4 top sgRNAs per gene for experiments. In our experience, when using nucleofection of RNPs, about 60-70% of such sgRNAs result in high editing efficiency (>80% indel efficiency). Total editing efficiency can usually also be increased by simultaneously nucleofecting with multiple sgRNAs targeting the same gene.

   b. If a near-precise deletion is desired, we typically design 2 sgRNAs spaced ~50-300bp apart. We have tested up to 1000bp spacing and this works well, although efficiencies for deletions <300bp can be more easily assessed (compared to larger deletions), via simple Sanger sequencing. When the goal is a gene KO, we usually design both sgRNAs to be in exons (or the same exon), so that even alleles that do not receive a deletion still have a good chance of resulting in gene KO via small indels.

   c. By necessity, available sgRNA design tools list possible sgRNA sequences based on a reference genome for the species you are using. It is a good idea to check the genomic sequence of your particular cells for polymorphisms (and mutations in the case of cancer cell lines) that may exist and may cause some sgRNA sequences created by design tools to have mismatches in your particular cells. In other words, it is best to ensure that the exact sgRNA sequence you decide to use is actually intact in your cells' genome.

2. CRISPR-RNP nucleofection:

*Note: Before starting, wipe down workspace and equipment with RNaseZap (Thermo Fisher) to inhibit RNases that may degrade your sgRNA. Wear clean gloves and use pipet tips designated for RNA work only.*

a. Reconstitute your synthetic RNA (and dilute it if desired):

   i) Briefly centrifuge your tubes or plates containing synthetic modified single guide RNA (sgRNA) oligos to ensure that the dried RNA pellet is collected at the bottom.

   ii) Carefully dissolve sgRNA in the provided nuclease-free 1X TE Buffer (Tris-EDTA, pH 8.0) by adding 15µl buffer to 1.5nmol of dried sgRNA. (This will result in a final concentration of 100µM (100pmol/µl)). Flick the bottom of the tube gently so the liquid covers the bottom, then pulse vortex the tube for ~30 seconds and quick spin reconstituted RNA down. Dissolved RNA should be stored at -20 °C when not in use. Under these conditions, RNA will be stable for at least one year.

   iii) If desired, to create a working stock, make a 30µM sgRNA dilution (e.g. 14µl nuclease-free water + 6µl of reconstituted 100µM sgRNA, creating a total of 20µl of 30µM (30pmol/µl) sgRNA). Use diluted sgRNA immediately or store at -20 °C for up to three months. Keep RNA on ice while in use. (If you will make larger master mixes of RNPs using the same sgRNA(s), it may not be necessary to dilute your sgRNA. sgRNAs are diluted primarily to allow for easier pipetting volumes.)

⇒ *Tip: If reduced editing efficiencies are experienced over time, this may be due to improper handling and/or storage of RNA. It is best to store RNA in 1X TE Buffer long-term and to dilute with water only the amount required in the short-term. Freeze-thawing of RNA should also be kept to a minimum.*

b. Dilute Cas9:

   i) Briefly centrifuge the stock Cas9 (Aldevron sNLS-SpCas9-sNLS, 61 µM) to ensure that all liquid is at the bottom. To make a working stock, make a 1:5 Cas9:PBS dilution (e.g. 10µl PBS (pH 7.4) + 2µl Cas9, creating 12µl of 10.17µM (10.17pmol/µl) Cas9). Keep diluted/undiluted Cas9 on ice. Only make the amount of diluted Cas9 that you will need at any given time since it is best not to freeze the diluted Cas9.

⇒ *Tip: It is best to dilute Cas9 only in PBS (pH 7.4), not directly in Nucleofector solution. We have found that concentrated Cas9 can precipitate when added directly to some buffers.*

⇒ *Tip: We have tested several commercial sources of purified Cas9 protein and have found that Aldevron's product provides the highest editing efficiency at the lowest dose. Other sources or in-lab purified Cas9 can also be used but may require a higher molar dose for equivalent editing efficiency. Note also that we have compared purified Cas9 proteins with three slightly different versions of nuclear localization signal (NLS) location: sNLS-SpCas9-sNLS, 2XNLS-*

*SpCas9-sNLS, and sNLS-SpCas9-2XNLS. We found that these three varieties performed identically in our cells.*

c. Assemble RNP complexes:

i) Create complete Nucleofector Solution by mixing SG Cell Line Solution (or other solution specific to your cell type) with the provided Supplement at a 4.5:1 ratio (e.g. for one nucleofection, combine 16.4µl Cell Line Solution + 3.6µl Supplement). Make a master mix for multiple nucleofections. (Once combined, the complete Nucleofector Solution is stable for three months.)

ii) Assemble RNP complexes in complete Nucleofector solution, adding reagents in the order shown below. The amounts below will allow for 15 pmol total RNPs to be nucleofected into one cell sample (with 10% extra volume), at an sgRNA:Cas9 ratio of 2:1. You may make a master mix for multiple nucleofections using identical RNPs.

| | |
|---|---|
| ul complete Nucleofector Solution | 19.28 µl |
| ul Synthego gRNA (**30 pmol/ul**) | 1.10 µl |
| ul Cas9 (**10.17 pmol/ul**) | 1.62 µl |
| Total | 22 µl |

*⇒ Tip: We have also tested higher sgRNA:Cas9 molar ratios up to 9:1 and have not found an increased ratio to result in significantly higher editing efficiency in NSCs and GSCs. However, it is possible that other cell types may benefit from an increased ratio, and this should be empirically determined. Similarly, the optimal total RNP dose should also be empirically determined for other cell types.*

iii) Mix by pipetting and briefly centrifuge the sample so all the liquid is at the bottom.

iv) Incubate RNPs for 10-20 minutes at room temperature and then keep on ice (or at 4°C) until shortly before use. **Immediately before using complexed RNPs to resuspend cells (see below), make sure to re-equilibrate RNPs back to room temperature.**

d. Collect and prepare cells:

i) Prepare a 12-well or 6-well dish (pre-coated if necessary, following your standard culturing procedure) so it contains appropriate culture media and pre-warm the plate by placing in a 37°C incubator until you are ready to plate your nucleofected cells. (The size dish that the cells will be plated into after nucleofection should be empirically determined so that your cells are ~30-50% confluent the day after nucleofection. This will vary depending on the size of your particular cells and how many die due to the nucleofection process.)

ii) Harvest cells normally and count. For each sample to be nucleofected, aliquot 1.5-2 x $10^5$ cells into a separate 1.5 mL microcentrifuge tube and centrifuge at 300g for 5 minutes.

iii) Resuspend cells in PBS to wash, and centrifuge at 300g for 7 minutes.

iv) Aspirate the liquid as completely as possible. Resuspend the cell pellet in 20μL of RNP complexes. Work quickly, but carefully, and avoid leaving cells in Nucleofector Solution for longer than 30 minutes total (from the time you resuspend them to the time nucleofection is complete). Avoid bubble formation.

v) Transfer all of the cell-RNP solution to one well of a 16-well Nucleocuvette strip (placed inside the supplied "plate skeleton" if using the 96-well Shuttle System), and cover with the provided lid (either strip lid or plate lid if using the Amaxa 4D Nucleofector X unit or the 96-well Shuttle Unit, respectively). Make sure there are no bubbles in your Nucleocuvette.

e. Nucleofect cells:

i) If using the Amaxa 4D Nucleofector X unit, turn on the core unit and then use the touch screen to select the appropriate options for nucleofecting a 16-well strip. Alternatively, if using the Amaxa 96-well Shuttle Unit, turn on the Amaxa core unit, then the 96-well Shuttle Unit, and then the laptop computer attached to the 96-well Shuttle Unit. Then open the 96-well Shuttle software.

ii) Visually inspect the Nucleocuvette Vessel to make sure that the sample covers the bottom of the cuvette and that there are no bubbles in the cuvette. If you notice problems, gently tap the Nucleocuvette Vessel against your hand and/or use a thin sterile pipet tip to pop any bubbles.

iii) If using a 16-well Nucleocuvette strip, insert the strip into the open Nucleofector 4D X unit. Make sure the larger gap in the strip lid is at the top of the strip rather than at the bottom, so that the yellow indicator in the X unit fits through the large gap at the top of the lid. Alternatively, if using the 96-well Shuttle Unit, place the Nucleocuvette Vessel with plate lid into the retainer of the 96-well Shuttle Unit. Check for proper orientation of the plate (A1 should be at the top left).

iv) Run nucleofection program EN-138 (or other cell-type specific program). After run completion, the screen should display a "+" over the wells that were successfully electroporated. Remove the cuvette strips/plate from the machine.

v) Carefully resuspend the cells in each well of the Nucleocuvette with 80μl of pre-warmed growth media, and mix very **slowly and gently** by pipetting up and down 3-4 times.

*⇒ Tip: Excessive pipetting can greatly increase cell death, so keep pipetting to a minimum. Furthermore, for some cell types Lonza recommends incubating the Nucleocuvettes at room temperature or 37°C for ~20 minutes immediately after nucleofection (before resuspending with media), as this can greatly increase cell survival. We have not found this to be necessary for NSCs and GSCs, but it may be important for other cell types.*

vi) Transfer all 100μl to one well of the pre-warmed, media-containing cell culture plate and gently pipet up and down a few times.

vii) Shake the plate to distribute the cells. Leave the plate at room temperature for ~20-

25 minutes and then place in incubator. (This helps distribute cells evenly.)

⇒ *Tip: Cell survival is usually highest if you keep the cells somewhat dense after nucleofection and do not plate them too sparse, i.e. plate them into an appropriate well/plate so they are ~30-50% confluent the day after nucleofection. Exact cell numbers to use and size well they should be plated into should be empirically determined for your particular cell type.*

f. Change the media 12-24 hours after nucleofection to remove dead cells and debris.

g. Three days after nucleofection, cells may be collected for indel analysis and/or phenotypic analysis, or cells may be expanded if creating a stable line. If studying an essential gene, phenotypic analysis may need to be done sooner as the majority of editing happens within 24 hours and the stability of the particular remaining protein will determine how quickly a phenotype can be observed.

⇒ *Tip: If absolutely necessary, cells can be passaged only 24 hours post nucleofection. However, this should be avoided if possible, as we have found that it results in reduced cell health and proliferation.*

3. <u>Editing efficiency analysis:</u>

a. Harvest nucleofected cells *and control cells* (nucleofected with RNPs containing a non-targeting sgRNA or mock nucleofected) according to normal culturing procedure. Cell pellet can either be used immediately or stored at -20°C or -80°C for several weeks before gDNA extraction.

b. Extract gDNA using a column purification kit, according to the manufacturer's protocol. We typically use the MicroElute Genomic DNA Kit (Omega Bio-Tek).

c. Perform PCR amplification of the region around the sgRNA cut site(s):

i) Whenever possible, design PCR primers to be ~250-350bp outside of the sgRNA cut site. If multiple sgRNAs are being used, design primers to be ~250bp outside of the outermost cut sites. We usually devise a ~1000bp (wt) amplicon. Use webtools such as Primer3 and NCBI Primer-Blast to check primer parameters and check for off-target amplification.

ii) Design one or multiple sequencing primers that are ~20-80bp inside either end of the PCR amplicon.

iii) Perform PCR amplification (50μl reaction) on the edited and control samples using a high-fidelity polymerase such as Phusion (New England BioLabs), following manufacturer's suggestions for PCR conditions for the particular enzyme. We typically include a final concentration of 3% DMSO (which aids with template denaturation) in our PCR reactions with gDNA templates. (See Table S1 for details on primer sequences, annealing temperatures, and extension times that we used for the sgRNAs described in our study.)

iv) Run 10μl of the PCR reactions on a 1% agarose gel and visualize with ethidium bromide staining. Check that there is only one product of the correct size in the control sample (the edited sample may have multiple bands if your RNPs created one or more deletions). If you have multiple products in the control sample, design new PCR primers and repeat the amplification.

⇒ *Tip: Sometimes it is simply not possible to identify PCR primers that only produce one band. If you have tried multiple primer sets and this is the case, you may be able to use a primer set that creates multiple bands. We have found that as long as the correctly-sized band is by far the most intense band, the following steps using simple column purification of the PCR product can still work properly, as long as a unique internal sequencing primer is used (a sequencing primer that does not overlap significantly with either PCR primer). Alternatively, if your best PCR primers produce multiple bands, you can also employ gel purification and only sequence a specific band. However, we do not recommend this since you may not sequence certain indels that are present in your edited sample if you only select a specific portion of the gel to sequence (rather than sequencing everything present in your PCR reaction).*

d. Use a column PCR purification kit to purify the remainder of the edited and control sample PCR products (~40μl), according to the manufacturer's protocol. We typically use the Monarch PCR & DNA Clean-up Kit (New England BioLabs).

e. Perform Sanger sequencing on the PCR amplicons for your edited and control samples.

f. Perform editing analysis using Sanger trace files:

i) Using a freely-available webtool such as Inference of CRISPR edits (ICE, https://ice.synthego.com/#/), enter the trace files (.ab1 files) for your edited and control samples, as well as the sequence(s) of the sgRNA(s) that you used. ICE can be used for samples where up to 3 sgRNAs were used simultaneously.

ii) ICE results will display information including your predicted total editing efficiency (called "ICE score"), predicted KO score (percent of predicted sequences that result in a frameshift or an indel ≥21 bp in length), and an $R^2$ value that describes "goodness of fit". Furthermore, the output allows visualization of indel size distribution, a list of the sequences predicted in your cell pool (in order of frequency of occurrence), and a discordance plot showing where (or if) your edited trace deviates from your control trace. Find more detailed explanation of outputs at https://www.synthego.com/guide/how-to-use-crispr/ice-analysis-guide.

**Supplies required for nucleofection:**

(does not include cell culture reagents)

| *Item* | *Company* | *Catalog number* |
|---|---|---|
| 2'-O-methyl 3'phosphorothioate-modified, chemically synthesized sgRNA | Synthego | N/A *** |
| 1X TE Buffer (Tris-EDTA, pH 8.0) | Synthego | Provided with sgRNA kit |
| Nuclease-free water | Synthego | Provided with sgRNA kit |
| sNLS-SpCas9-sNLS nuclease | Aldevron | 9212-0.25MG |
| Phosphate buffered saline, pH 7.4 | homemade or e.g. Fisher | e.g. 10010-023 |
| SG Cell Line 4D-Nucleofector™ Kit (20µl format) | Lonza | V4XC-3032 (32 reactions)<br>V4SC-3096 (96 reactions) |

****For this protocol we necessarily used Lonza's instruction manual for the Amaxa 4D Nucleofector as a reference. See also Synthego's CRISPR RNP nucleofection protocol.*
